# Ongoing evolution of Middle East Respiratory Syndrome Coronavirus, Kingdom of Saudi Arabia, 2023-2024

**DOI:** 10.1101/2024.09.12.612455

**Authors:** Ahmed M. Hassan, Barbara Mühlemann, Tagreed L. Al-Subhi, Jordi Rodon, Sherif A. El-Kafrawy, Ziad Memish, Julia Melchert, Tobias Bleicker, Tiina Mauno, Stanley Perlman, Alimuddin Zumla, Terry C. Jones, Marcel A. Müller, Victor M. Corman, Christian Drosten, Esam I. Azhar

## Abstract

Middle East respiratory syndrome coronavirus (MERS-CoV) circulates in dromedary camels in the Arabian Peninsula and occasionally causes spillover infections in humans. Due to lack of sampling during the SARS-CoV-2 pandemic, current MERS-CoV diversity is poorly understood. Of 558 dromedary camel nasal swabs from Saudi Arabia, sampled November 2023 to January 2024, 39% were positive for MERS-CoV RNA by RT-PCR. We generated 42 MERS-CoV and seven human 229E-related CoV by high-throughput sequencing. For both viruses, the sequences fell into monophyletic clades apical to the most recent available genomes. The MERS-CoV sequences were most similar to those from lineage B5. The new MERS-CoVs sequences harbor unique genetic features, including novel amino acid polymorphisms in the Spike protein. The new variants require further phenotypic characterization to understand their impact. Ongoing MERS-CoV spillovers into humans pose significant public health concerns, emphasizing the need for continued surveillance and phenotypic studies.

## Introduction

Middle East respiratory syndrome coronavirus (MERS-CoV) was first described in 2012 in a single human case of viral pneumonia.^1^ Subsequent research uncovered a widespread zoonotic disease caused by a virus that infects humans in countries of the Middle East as well as East and West Africa.^2–7^ The virus is primarily acquired via direct contact with dromedary camels, the main reservoir host, and with lesser efficiency through contact with infected humans.^8–13^ Human-to-human transmission in household and community settings is limited, but nosocomial outbreaks with inter-hospital transmission and prolonged transmission chains have occurred.^14–17^ Even subtle changes in viral transmissibility may thus enable the virus to adapt to humans and spread in the manner of an epidemic or pandemic. Due to the zoonotic nature of MERS-CoV, virus evolution in dromedary camel populations is of immediate relevance to human disease.^18^

MERS-CoV can be differentiated into three distinct clades (A, B, and C). Clade A and B viruses are associated with dromedary camels in the Arabian Peninsula, clade C viruses with African dromedary camels. Clade A viruses have not been detected after 2015 and may be extinct. Clade B viruses have been circulating and evolving in Arabian dromedary camels at least up to 2019^19^ and have been classified into five phylogenetic lineages.^20^ While there is evidence for limited infectivity and virulence associated with clade C viruses derived from African dromedary camels, viruses of lineage B5 show signs of increased virulence and fitness in both dromedary camels and humans.^17,21–25^

In addition to MERS-CoV, dromedary camels also harbor a coronavirus closely related to seasonal human coronavirus (HCoV) 229E (subgenus *Duvinacovirus*),^26,27^ highlighting the importance of dromedary camels as a reservoir host for coronaviruses.

Due to a lack of sampling during the SARS-CoV-2 pandemic, there is limited knowledge regarding the diversity of currently circulating MERS-CoVs in the Arabian Peninsula.^28^ It remains unknown whether lineage B5 continues to dominate in camel populations of the Arabian Peninsula, as it did between 2017 and 2019, and whether currently circulating MERS-CoVs carry polymorphisms that might impact transmissibility or virulence. From January to May 2024, four laboratory-confirmed cases were reported to the World Health Organization by the Ministry of Health of Saudi Arabia (https://www.who.int/emergencies/disease-outbreak-news/item/2024-DON516, accessed 2024-05-31), indicating continuous zoonotic spillover into the human population. There is a need for virus surveillance to monitor ongoing changes in the viral genomes. Here we present the genetic characterization of 42 MERS-CoV genomes from infected dromedary camels sampled in KSA from late November 2023 to early January 2024.

## Results

Between November 2023 and January 2024, during the dromedary camel breeding period, 558 dromedary camels were sampled in camel farms in six locations in Saudi Arabia (Jeddah: n=101, Al Duwadimi: n=90, Al Quwayiyah: n=97, Al Riyadh: n=108, Sajir: n=94, Shaqra: n=68). A total of 217 were found to be positive for MERS-CoV RNA by RT-PCR (Jeddah: n=36/101 (35·6%), Al Duwadimi: n=63/90 (70·0%), Al Quwayiyah: n=21/97 (21·6%), Al Riyadh: n=58/108 (53·7%), Sajir: n=28/94 (29·8%), Shaqra: n=11/68 (16·2%)), corresponding to an overall positivity rate of 38.9% (n=217/558) (Table 1, Table S1). We selected 51 samples for sequencing based on viral load >2.10×10^6^ genome copies/ml, resulting in 42 MERS-CoV genomes with >99% genome coverage and coverage depth >5x. Samples yielding MERS-CoV sequences were collected at six sampling sites between 2023-11-22 and 2024-01-05 (Jeddah: n=6, Al Duwadimi: n=4, Al Quwayiyah: n=8, Al Riyadh: n=8, Sajir: n=12, Shaqra: n=4). Twenty-two samples were found to harbor a co-infection with a HCoV-229E-related CoV, resulting in 7 near-complete genomes (>94% genome coverage, Jeddah: n=5, Sajir: n=2, Table 1, Table S1).

**Table 1:**
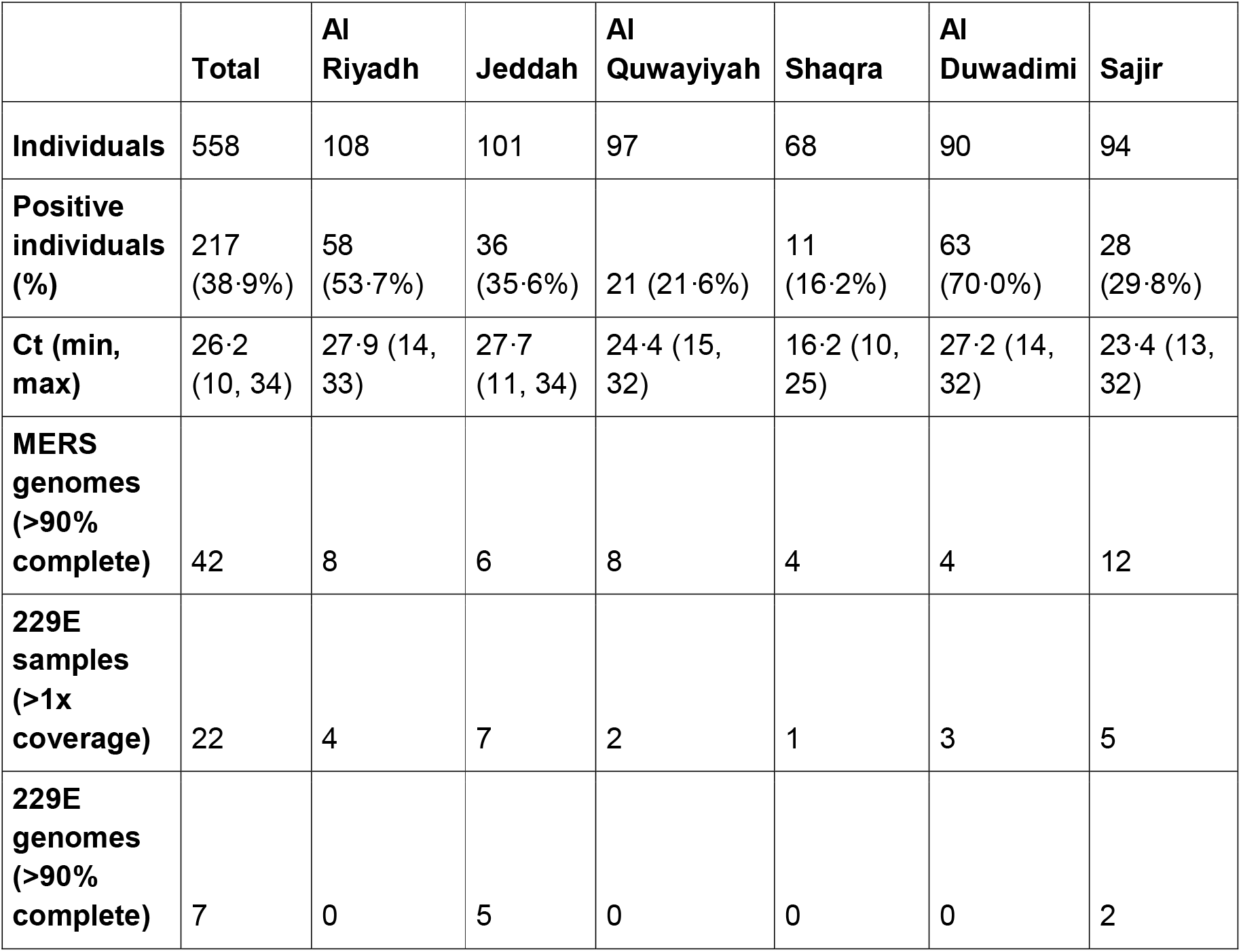
Number of positive samples across sampling sites.

To investigate the presence of mixed infections or laboratory contamination, we considered all genome positions where the majority base was present in <80% of all reads at positions with >5x coverage. The median number of such minor variant positions per sample was 1 (range: 0-118, fig. S1). The sample with 118 minor variant positions was excluded from further phylogenetic analyses. The remaining MERS-CoV sequences generated in this study form a monophyletic clade apical to lineage B5, here called B5-2023 (Fig. 1A). The B5-2023 clade is part of a ladder-like pattern of evolution within lineage B5 (Fig. 1A). Based on a phylogenetic tree inferred from the full genome (Fig. 1A,B) five sub-lineages within the B5-2023 clade can be differentiated (B5-2023.1 - B5-2023.5). Recombination analysis detected a previously described recombination event between lineages B3 and B4 preceding the formation of lineage B5,^26^ which was also found in the sequences generated here (Fig. S2A,B). Furthermore, we found indications of three recombination events within the monophyletic B5-2023 clade (Fig. S2): First, a recombination event between sequences from clade B5-2023.1 and clade B5-2023.4 with breakpoints at positions 17,816 and 29,588, may have resulted in sequence ‘Al Quwayiyah/F6-P4b/B5-2023.1’ (Fig. S2C,D). Second, sequences of clade B5-2023.3 may have arisen from a recombination event between sequences from clade B5-2023.2 and B5-2023.4 with breakpoints at positions 751 and 15,585 (Fig. S2E,F). Third, the phylogenetic trees in Fig S2D,F also suggest that sequence ‘Al Duwadimi/P6-25/B5-2023.5’ may have arisen as the result of a recombination event between a lineage B5 sequence basal to the B5-2023 cluster and sequences from clade B5-2023.3 (Fig. S2G).

**Figure 1:**
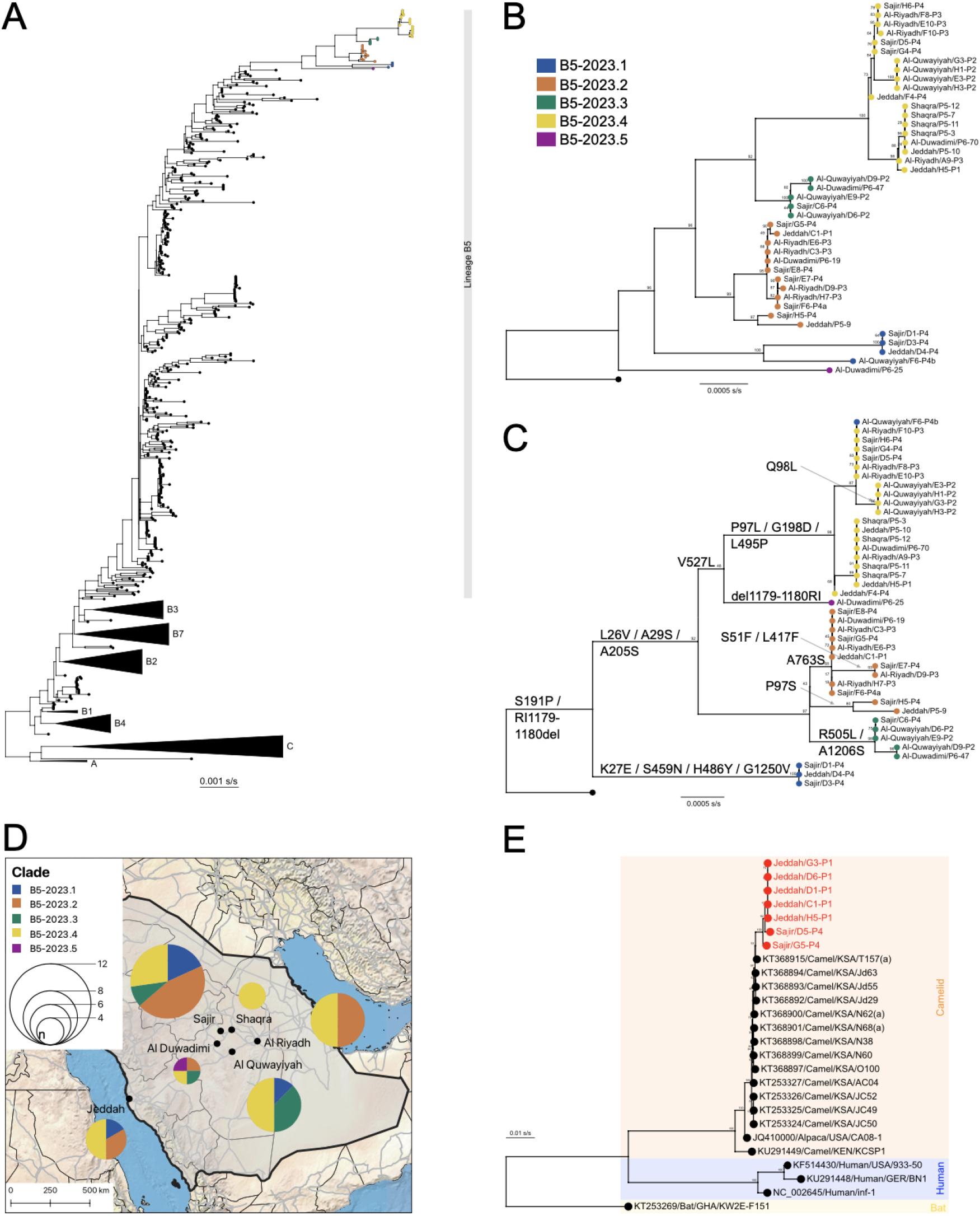
Phylogenetic trees and distribution of samples. A) Maximum likelihood phylogenetic tree of 620 complete MERS-CoV-genomes sampled until 2019 (black tips), and 41 genomes generated in this study (colored tips). Clades A, and C, as well as lineages B1-B4 and B7 are collapsed. B) Maximum likelihood tree of clade B5-2023 sequences generated in this study. The tree is rooted with a lineage B5 sequence from 2019 (GenBank accession No. OL622036.1) and numbers on nodes show bootstrap support. Sequences are separated into sub-clades based on monophyly (blue: B5-2023.1, orange: B5-2023.2, green: B5-2023.3, yellow: B5-2023.4, magenta: B5-2023.5). C) Maximum likelihood tree of Spike sequences of clade B5-2023. Amino acid substitutions are annotated. The same coloring and root is used as in panel B. D) Spatial distribution of sub-lineages within the B5-2023 clade. Pie charts show the number of sequences from each sub-clade found at each sampling site, with the size of the circle corresponding to the number of sequences. Administrative regions are delineated by light black lines, roads by light-gray lines. E) Maximum likelihood phylogenetic tree of 26 HCoV-229E-related CoV sequences. Sequences from this study are highlighted in red. Stars show nodes with bootstrap support higher than 70. The tree was rooted with KT253269/Bat/GHA/KW2E-F151. KSA: Kingdom of Saudi Arabia, KEN: Kenya, USA: United States of America, GER: Germany, GHA: Ghana, s/s: substitutions/site.

A regression of root-to-tip distances against sampling dates suggests a constant clock rate across the entire tree as well as for lineage B5 and the B5-2023 clade (Fig. S3A,B). The sequences in the B5-2023 clade have acquired 39 nucleotide substitutions compared to previous lineage B5 sequences circulating until 2019 (Table S2, S3). These polymorphisms include a substitution (S191P) and a two amino acid deletion (R1179-I1180del) in the N-terminal and S2 domains of the Spike protein, respectively. Sequences within the B5-2023 clade have acquired a total of 21 substitutions in the Spike protein across the whole clade (Fig. 1C, Table S2), including substitutions in the receptor binding domain (RBD) (S459N, H486Y, L495P, R505L, V527L) and within the cathepsin L cleavage site (A763S).^29^ The sequences did not have any deletions of accessory open reading frames, as often found in clade C viruses.^25,30,31^ We did not find evidence for geographic clustering of sub-lineages within the new clade, with five of the six sampling sites showing circulation of at least two sub-lineages and all but one sub-lineage being detected in at least three of the six sampling sites (Fig. 1D).

The seven *Duvinacovirus* sequences formed a monophyletic clade apical to previously detected HCoV-229E-related CoV from dromedary camels (Fig. 1E) sampled in 2014/2015. Regression of root-to-tip distances and sampling dates (Fig. S3C) shows a constant clock rate in the data. No deletions in ORF8, as described by Corman et al.,^27^ were found.

## Discussion

The present study identifies a novel monophyletic clade of MERS-CoVs currently circulating in Saudi Arabia. While previously circulating lineages within clade B arose from deep splits in the phylogenetic tree, the apical branching of the novel sequences from lineage B5, as well as the ladder-like tree topology of lineage B5, suggest that present MERS-CoV strains in the Arabian Peninsula originated from the sole circulation of lineage B5. This is also supported by the recombination analysis, and the indications of a constant molecular clock rate shown in this study.

Twenty-one different substitutions in the Spike protein are present in the MERS-CoV sequences in the B5-2023 clade (Figure 1C). A number of those are located in the RBD or NTD (Table S2). The RBD has been shown to be the site most frequently targeted by neutralizing antibodies.^32^ Two of the substitutions found in the B5-2023 clade (L495P and V527L) are situated on the ridge of the receptor binding motif exposed in the closed conformation of the Spike, an epitope preferentially targeted by antibodies in human post-infection sera.^32,33^ Furthermore, a substitution in the cathepsin L cleavage site (A763S) present in clade B5-2023.3, may affect Spike cleavage and virus infection in cells that do not express TMPRSS-2.^29^ These substitutions require further study to investigate their phenotypic effects on virus entry, receptor affinity, immune escape, and replicative fitness.

Among the diversity of currently circulating MERS-CoVs, the distinct sub-lineages did not cluster geographically, showing that dromedary reservoirs are maintaining virus diversity across the Arabian Peninsula. Given the reservoir traits of dromedary camels, including rapid viral clearance, waning adaptive immune responses and evidence of rapid re-infection,^34–38^ it is very likely that parallel evolution of distinct MERS-CoV sub lineages is ongoing in dromedary camels. This is consistent with pre-2020 studies of MERS-CoV genetic diversity^39,40^ and with the detection of three recombination events within the B5-2023 clade. The movement of camels for grazing and leisure results in mixing of populations from different geographical regions,^41^ which might facilitate MERS-CoV spread across the Arabian Peninsula. In-depth epidemiological and spatiotemporal studies are urgently needed to identify hotspots for MERS-CoV dissemination and areas with high risk of human spillover.

Forty-three percent of the 51 sequenced samples showed evidence of infection with HCoV-229E-related CoV. Widespread infection of dromedary camels with that virus, both with and without MERS-CoV co-infection, has been observed previously.^26,27^ The apical placement of the newly described HCoV-229E-related sequences, together with temporal signal observed here, may also point to a ladder-like pattern of evolution of the virus in dromedary camels.

Similarities in the epidemiology of HCoV-229E-related CoV and MERS-CoV in dromedary camels, including the absence of severe disease and the higher rate of infection in younger camels,^27^ suggest that HCoV-229E-related CoV may be maintained at population level in a similar fashion as MERS-CoV, highlighting the importance of dromedary camels as reservoir hosts for coronaviruses.

Ongoing spillovers of MERS-CoV into the human population in the Arabian Peninsula pose a significant public health concern. Global efforts to mitigate MERS-CoV impact in human populations are threatened by lineage B5 viruses, which possess enhanced replicative fitness and transmission capabilities in dromedary camels.^21–24^ The MERS-CoV genome evolution revealed by the sequences presented here, the first available from the Arabian Peninsula since 2019, highlight the urgent need for further MERS-CoV surveillance and phenotypic studies.

## Acknowledgements

We thank Nikolai Zaki and Annowah El-Duah for technical assistance.

This work was funded by the German Federal Ministry of Education and Research through project DZIF (8040701710 and 8064701703) and VARIpath (01KI2021), the Federal Ministry of Health through Grant SeroVarCoV, EU Hera project DURABLE (101102733), as well as the Jameel Fund for Infectious Diseases Research and Innovation.

MAM receives funding from DFG (MU 3564/3-1) and NSF (no. 125599).

CD received additional funding from ECDC project Aurorae (NP/21/2021/DPR/25121).

VMC is a participant in the BIH-Charité Clinician Scientist Program funded by Charité - Universitätsmedizin Berlin and the Berlin Institute of Health.

VMC and MAM have their names on patents regarding SARS-CoV-2 serological testing and monoclonal antibodies.

## Author contributions

Ahmed M. Hassan: Validation, Investigation, Resources, Data curation, Writing - Review and editing

Barbara Mühlemann: Software, Validation, Formal analysis, Investigation, Data curation, Writing - Original draft, Writing - Review and editing, Visualization

Tagreed L. Al-Subhi: Resources, Writing - Review and editing

Jordi Rodon: Investigation, Writing - Original draft, Writing - Review and editing Sherif A. El-Kafrawy: Resources, Writing - Review and editing

Ziad Memish: Resources, Writing - Review and editing

Julia Melchert: Methodology, Investigation, Writing - Original draft, Writing - Review and editing

Tobias Bleicker: Methodology, Validation, Investigation, Writing - Review and editing

Tiina Mauno: Methodology, Validation, Investigation, Writing - Review and editing

Stanley Perlman: Writing - Review and editing

Alimuddin Zumla: Resources, Writing - Review and editing

Terry C. Jones: Software, Writing - Review and editing, Supervision

Marcel A. Müller: Writing - Review and editing, Supervision, Funding acquisition

Victor M. Corman: Writing - Review and editing, Supervision, Funding acquisition

Christian Drosten: Conceptualization, Writing - Original draft, Writing - Review and editing, Supervision, Funding acquisition

Esam I. Azhar: Conceptualization, Resources, Writing - Review and editing, Supervision, Funding acquisition

