## Supplementary material for "Ongoing evolution of Middle East Respiratory Syndrome Coronavirus, Kingdom of Saudi Arabia, 2023-2024"

### **Methods**

#### **Sample collection**

A total of 572 nasal swabs samples taken from 558 individuals were collected from local camel farms in Saudi Arabia. Local camels were sampled from local farms in Jeddah (western Saudi), Al Quwaiiyah, Shaqra, Sajir, Al Duwadimi, and Al Riyadh (all central Saudi). All swabs were collected, immersed in viral transport media (VTM), transported in a cold container, and stored at -80˚C until further analysis. Samples were collected upon ethical approval from the Unit of Biomedical Ethics in King Abdulaziz University Hospital.

#### **RNA-extraction and reverse transcription-PCR-screening**

Viral RNA was extracted from 200μl of the VTM sample using QiaAmp viral RNA extraction kit (Qiagen, Germany) according to the manufacturer’s instructions. Testing for MERS-CoV was done using real-time reverse transcription-PCR (RT-PCR) assays targeting upE and ORF1A as described previously.[^42^](https://app.readcube.com/library/fefdf68c-7648-4f43-ae8d-20ef8d719102/all?uuid=07968020686720911&item_ids=fefdf68c-7648-4f43-ae8d-20ef8d719102:5625d20b-121a-4139-8f0c-76e269ff7a07) Samples that were positive for both targets with cycle thresholds (Ct) values <40 were considered positive.[^42^](https://app.readcube.com/library/fefdf68c-7648-4f43-ae8d-20ef8d719102/all?uuid=5801973966084555&item_ids=fefdf68c-7648-4f43-ae8d-20ef8d719102:5625d20b-121a-4139-8f0c-76e269ff7a07)

#### **Sequencing**

Full genomes were generated using shotgun Illumina High-Throughput Sequencing (HTS) on all selected positive samples and subsequent targeted enrichment was carried out if necessary.

HTS library preparation was performed using the KAPA RNA Hyper Prep Kit (Roche) according to the manufacturer's instructions. Briefly, 5 µL RNA were used for fragmentation at 85 °C for 6 min. Indexed DNA libraries were measured by Qubit dsDNA HS Assay kit (Thermo Fisher Scientific™, Massachusetts, USA) and Agilent TapeStation using the HS D1000 Kit (Agilent, California, USA). Equimolar pooled libraries were paired-end sequenced on a NovaSeq 6000 (paired end, 200 cycles, Illumina, California, USA).

In order to generate full-genome sequences, we applied a targeted enrichment approach using myBaits® (Daicel Arbor Biosciences, Ann Arbor, US). Here, we designed a capture bait-set using an alignment of 119 viral sequences, including reference sequences from MERS-CoV (n=20), SARS-CoV (n=39), SARS-CoV-2 (n=1), and the human endemic CoVs: OC43 (n=20), NL63 (n=15), 229E (n=10), HKU1 (n=5). The final bait-set comprised a total of 38,279 baits with a length of 80 nucleotides and three-fold tiling density. Among the generated baits, there were no BLAST hits to the following genomes: human, *Sus scrofa*, *Camelus dromedarius*, and *Myotis lucifugus*. The bait-set can be ordered from BioCat (Heidelberg, Germany) using our initial reference number: #200408-94. Targeted enrichment was performed as previously described for other viruses,[^43–45^](https://app.readcube.com/library/fefdf68c-7648-4f43-ae8d-20ef8d719102/all?uuid=514489696752553&item_ids=fefdf68c-7648-4f43-ae8d-20ef8d719102:58cf4195-b412-4ffd-9232-dee0d1640ea1,fefdf68c-7648-4f43-ae8d-20ef8d719102:fb08cdf9-c7f3-4eb9-ab3b-a941d723c12d,fefdf68c-7648-4f43-ae8d-20ef8d719102:b010b52c-572a-4c9a-be4c-2a85307e0cfb) following the manufacturer's recommendations. Hybridization was performed for 18 h at 65 °C; washing steps were carried out at 65 °C. The enriched libraries were amplified for 14 cycles using the KAPA Hifi HotStart Ready Mix and Library Amplification Primer Mix (Roche). Purified and quantified libraries were equimolar pooled and paired-end sequenced on a MiniSeq (150 cycles, Illumina).

#### **Bioinformatic analyses**

Next generation sequencing reads were trimmed using AdapterRemoval (version 2.3.2)[^46^](https://app.readcube.com/library/fefdf68c-7648-4f43-ae8d-20ef8d719102/all?uuid=03867083681455041&item_ids=fefdf68c-7648-4f43-ae8d-20ef8d719102:c2c38b61-6738-44b9-ba87-ac5423c1b1af) with the --qualitymax 41 --trimns --minlength 30 --trimqualities --minquality 2 options. Reads were mapped using kraken2 and the resulting krona plots were inspected for evidence of infection with HCoV 229E. Reads were mapped against OL622036.1 (MERS-CoV) and KT253327.1 (HCoV-229E-related CoV) with bowtie2 (version: 2.4.2)[^47^](https://app.readcube.com/library/fefdf68c-7648-4f43-ae8d-20ef8d719102/all?uuid=5972025406700069&item_ids=fefdf68c-7648-4f43-ae8d-20ef8d719102:60c668c2-9305-48f2-ac68-9458eb36ea8b) using the --no-unal --local options, reporting only the best match for each read. BAM files from the native and capture sequencing were merged using samtools. The consensus sequences were called using iVar (version: 1.3.1)[^48^](https://app.readcube.com/library/fefdf68c-7648-4f43-ae8d-20ef8d719102/all?uuid=8129444078000316&item_ids=fefdf68c-7648-4f43-ae8d-20ef8d719102:ff37238f-0696-46c4-bfdd-2d00aa1d8e4f) with the -m 5 -t 0.6 options.

All samples were screened for minor variants using code implemented in <https://github.com/VirologyCharite/minor-variants/>. A position was assumed to be a minor variant if the frequency of the most common nucleotide at that position was <80%, and the position was covered by more than 5 reads. The median number of minor variant sites per sample was 1 (range: 0, 118).

For phylogenetic analysis, all MERS-CoV genomes available under txid1335626 were downloaded from NCBI (2024-02-29) with the following search query: txid1335626[Organism:exp]. Sequences were filtered to include sequences longer than 29,900 nt and with >90% coverage, and excluding sequences of bat origin or low quality. This resulted in 620 genomes. The sequences were aligned with mafft (version: v7.471)[^49^](https://app.readcube.com/library/fefdf68c-7648-4f43-ae8d-20ef8d719102/all?uuid=004734535912886084&item_ids=fefdf68c-7648-4f43-ae8d-20ef8d719102:3d56431d-4f3e-4119-a3ff-58a9dff3fbf2) using the --auto --addfragments option, with the NC_019843 sequence as the reference.

Recombination analysis was performed in RDP4,[^50^](https://app.readcube.com/library/fefdf68c-7648-4f43-ae8d-20ef8d719102/all?uuid=255084065426457&item_ids=fefdf68c-7648-4f43-ae8d-20ef8d719102:2e3e1d7c-e4d7-4bab-a7e5-21ecdd4ff3da) including the sequences generated here as well as the lineage B5 sequences. The automated exploratory analysis implemented in RDP4 uses seven recombination detection algorithms (RDP, GENECONV, Chimaera, MaxChi, BootScan, SiScan, and 3Seq). The analysis identified two recombination events that involved any of the novel sequences, which were confirmed by inferring maximum likelihood trees from the minor and major parent in using iqtree (using the -m GTR+F+R4 -B 1000 -nt AUTO options, performing 1000 ultrafast bootstraps). Visual inspections of plots of pairwise sequence identity of sequences from lineages B3, B4, B5, and the novel clade described here, showed the presence of a similar recombination event between lineages B3 and B4 that formed lineage B5 also in the novel sequences described here. Finally, comparisons of trees made from the full genome, and trees made from the spike protein suggest an additional recombination event involving sequence ‘Al Duwadimi/P6-25/B5-2023.5’, possibly by parents involving a late 2019 lineage B5 sequence, and an B5-2023.3 sequence. Pairwise sequence identity was calculated for sequences excluding invariant sites, using a window size of 15 and a step size of 1.

Maximum likelihood phylogenetic trees were inferred using iqtree (version 2.2.0.3).[^51^](https://app.readcube.com/library/fefdf68c-7648-4f43-ae8d-20ef8d719102/all?uuid=9365846331313253&item_ids=fefdf68c-7648-4f43-ae8d-20ef8d719102:f6792069-29fa-4f09-8b99-a4415ad23ae6) The substitution model was determined by performing an initial iqtree run on the full alignments with the -m MF -nt AUTO options.[^52^](https://app.readcube.com/library/fefdf68c-7648-4f43-ae8d-20ef8d719102/all?uuid=8334335225271009&item_ids=fefdf68c-7648-4f43-ae8d-20ef8d719102:2a9719be-49ea-4ae0-a053-6e9597eb1d34) This determined the GTR+F+R4 substitution model as optimal for the MERS-CoV and HCoV-229E alignments. Subsequent iqtree runs were performed using the -m GTR+F+R4 -B 1000 -nt AUTO options, performing 1000 ultrafast bootstraps. Regression of root-to-tip distances against sampling dates was performed in TreeTime (version 0.11.3).[^53^](https://app.readcube.com/library/fefdf68c-7648-4f43-ae8d-20ef8d719102/all?uuid=05368609077086606&item_ids=fefdf68c-7648-4f43-ae8d-20ef8d719102:43d6f176-7bb9-47b9-9786-ea23d669c216) The trees from the previous iqtree runs were used as input for the full MERS-CoV and HCoV-229E trees. To investigate the temporal signal in lineage B5, a separate tree was inferred as described above, using just the MERS-CoV lineage B5 sequences and the novel sequences described here.

Figures were plotted in Python3.10 using the matplotlib,[^54^](https://app.readcube.com/library/fefdf68c-7648-4f43-ae8d-20ef8d719102/all?uuid=0873282585694577&item_ids=fefdf68c-7648-4f43-ae8d-20ef8d719102:3e6511b9-244a-425b-8971-e83aed8e9fa1) seaborn,[^55^](https://app.readcube.com/library/fefdf68c-7648-4f43-ae8d-20ef8d719102/all?uuid=9603484809163315&item_ids=fefdf68c-7648-4f43-ae8d-20ef8d719102:d845d84e-6d5e-4d3f-8238-c03607f07654) and baltic (<https://github.com/evogytis/baltic>) packages. The map was generated using QGIS (version 3.28).

### **Figures**

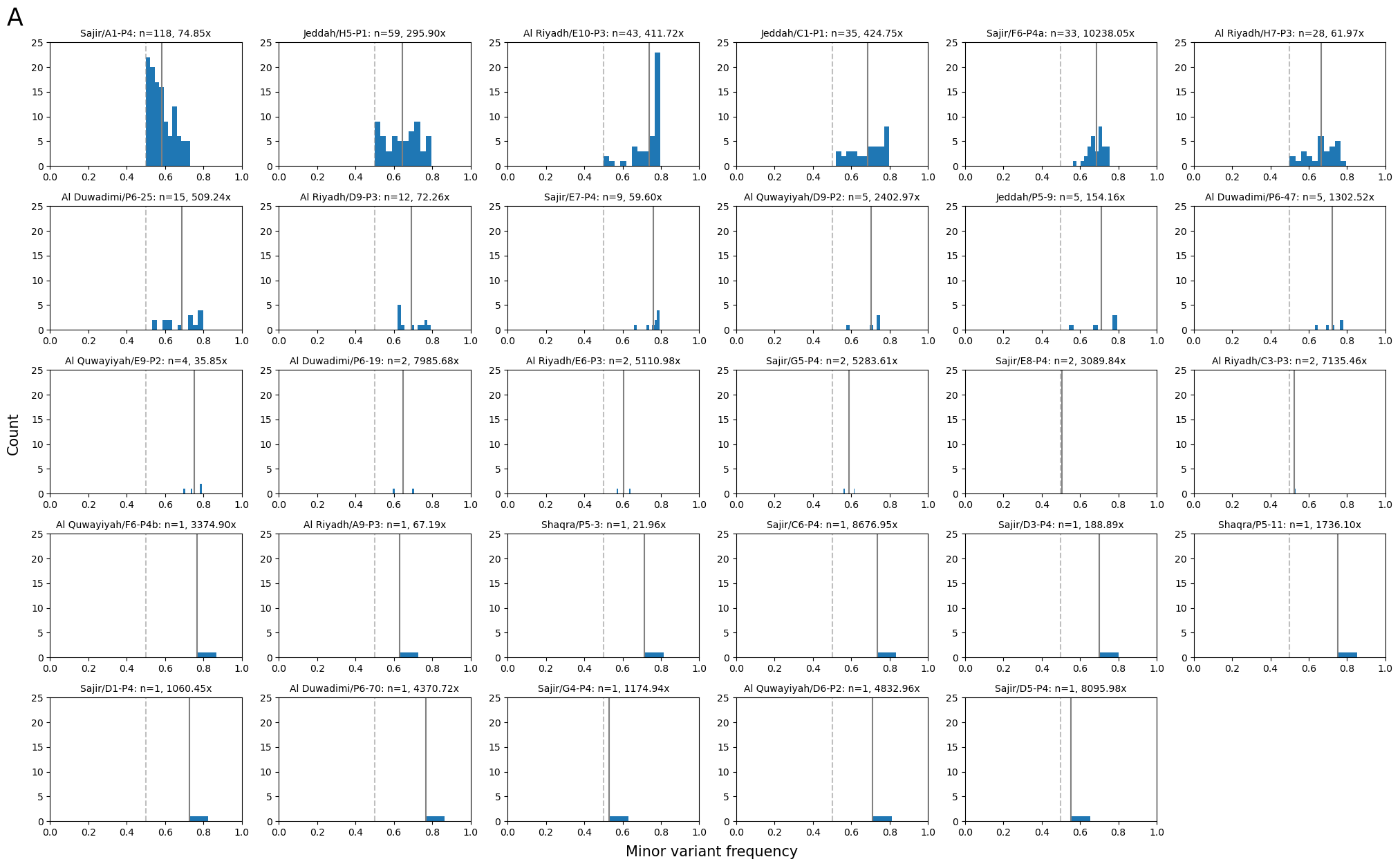

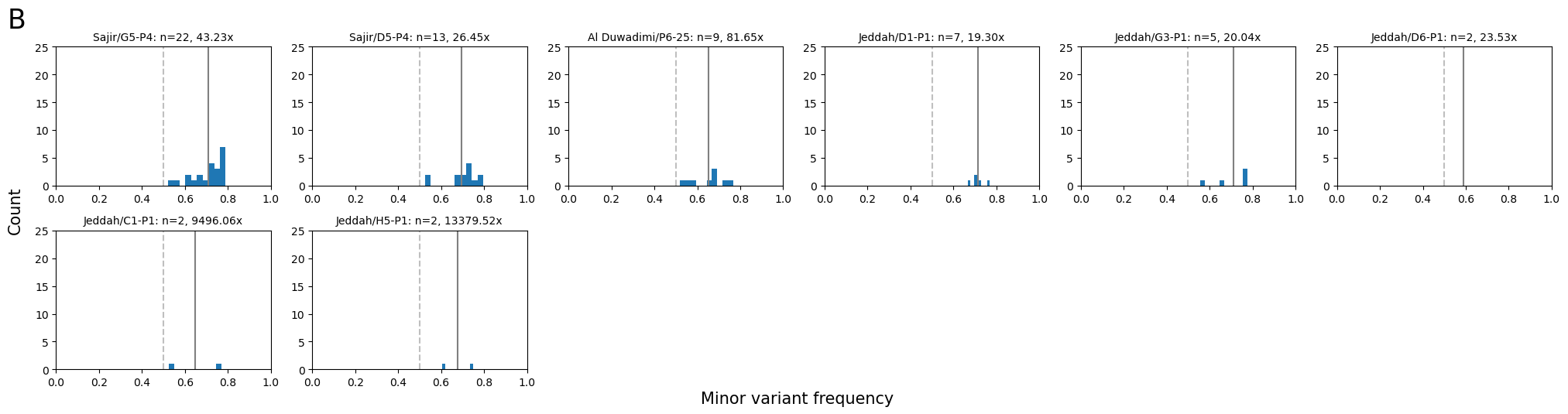

**Figure S1: Distribution of the frequencies of the most common base at minor variant sites.** A) MERS-CoV, B) HCoV-229E-related CoV. A site is defined as a minor variant site if it is covered by at least five reads and the frequency of the most common base is less than 0.8. Bold lines show the mean, dashed lines show 0·5 minor variant frequency. The title indicates the sample name, the number of minor variant sites, and the mean coverage depth of the entire sample.

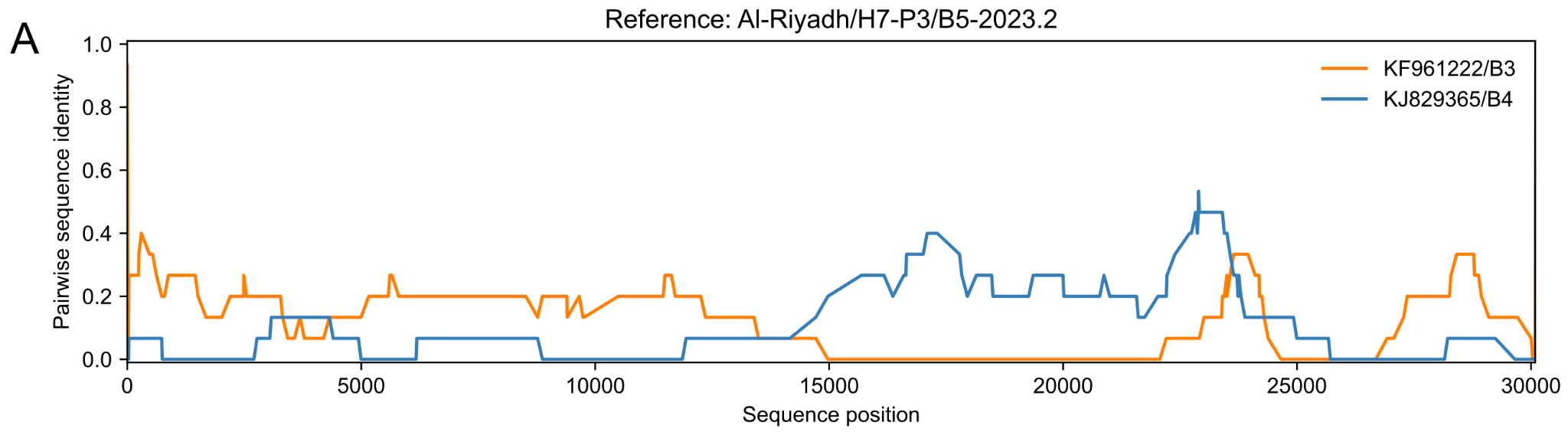

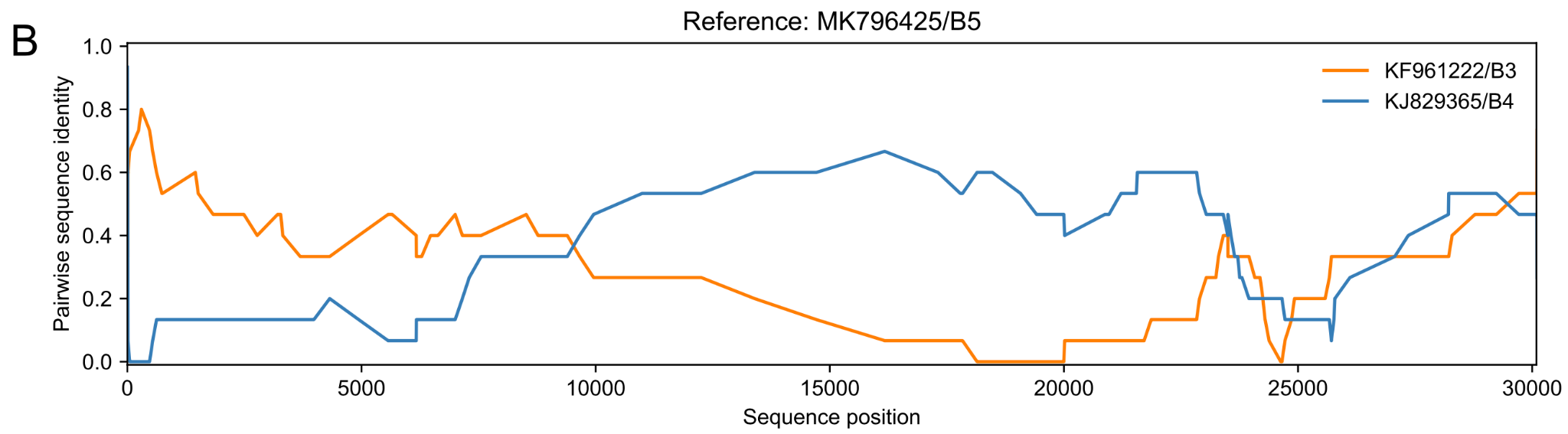

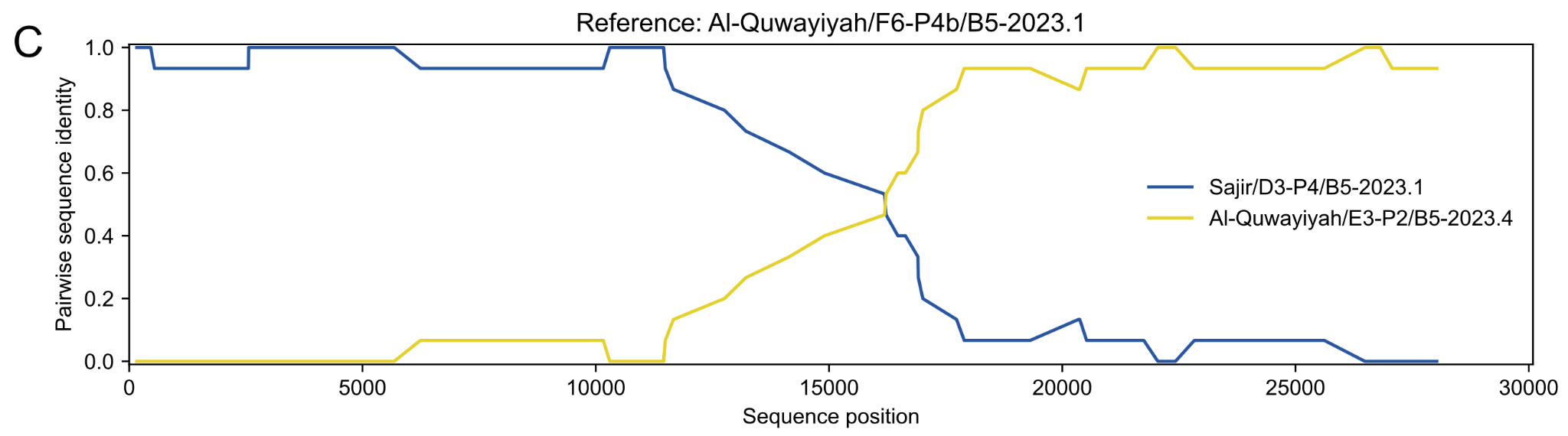

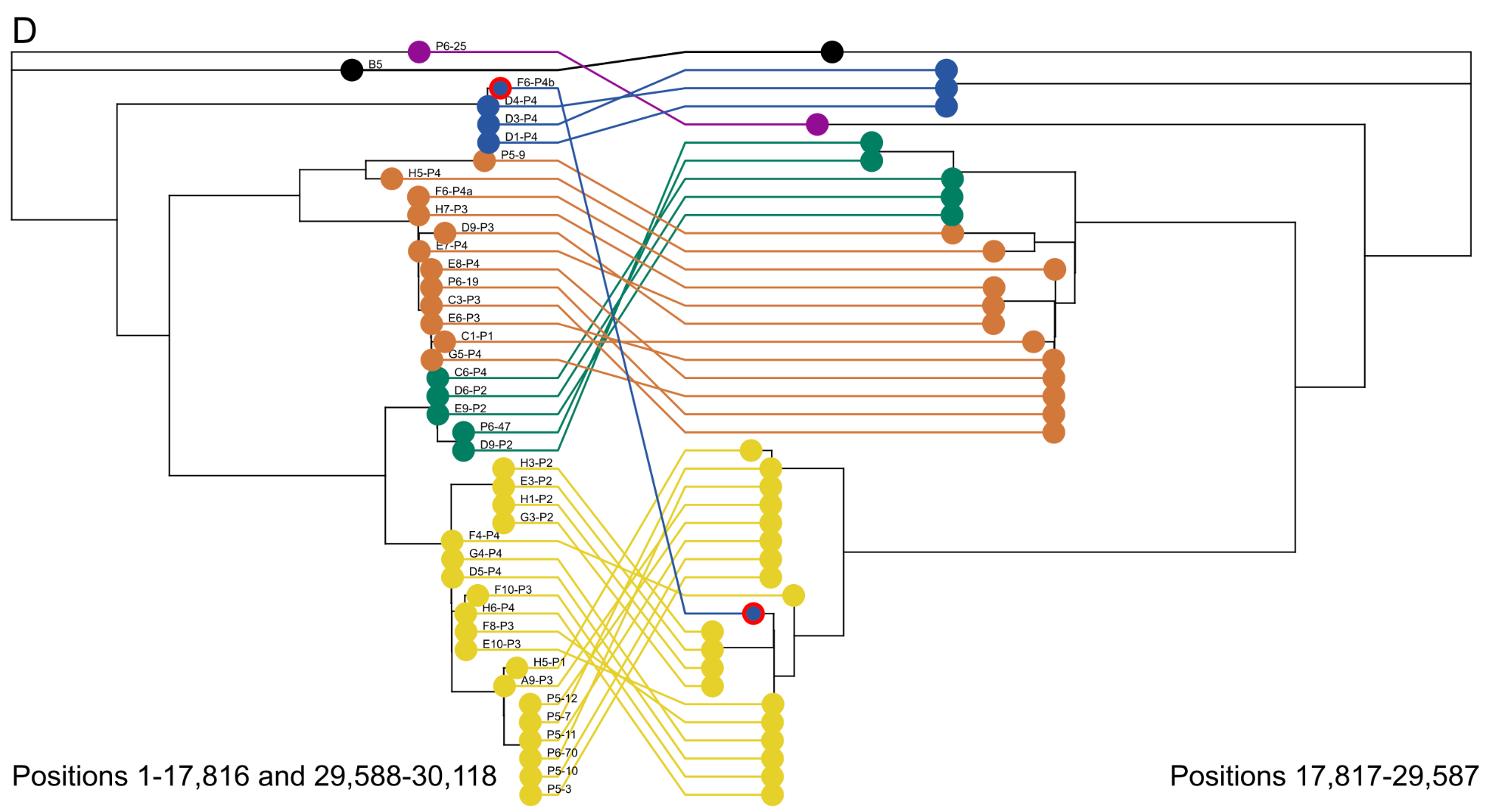

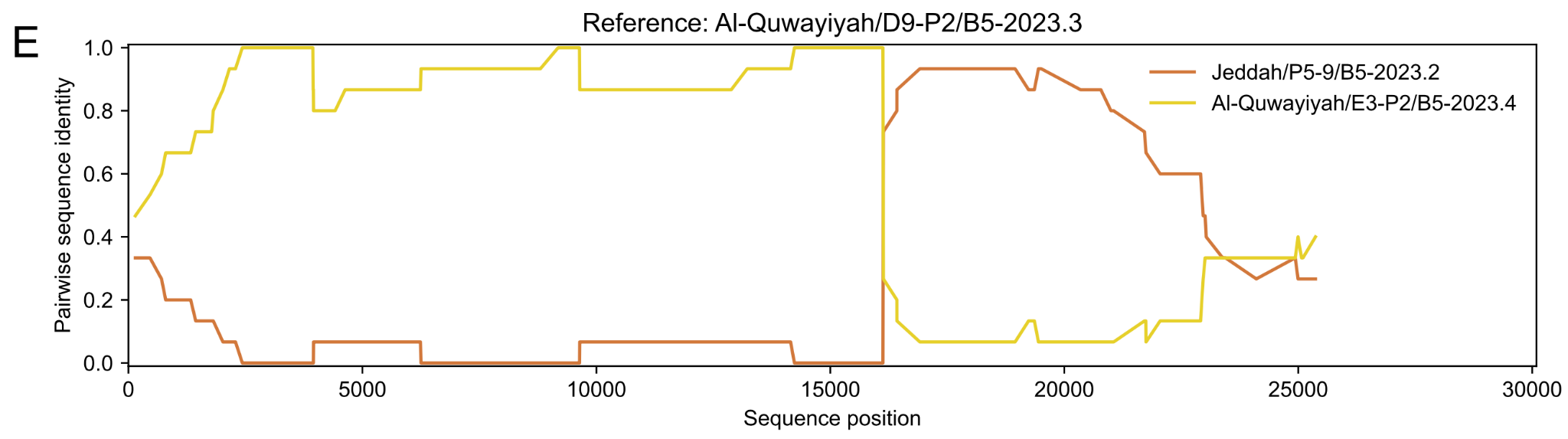

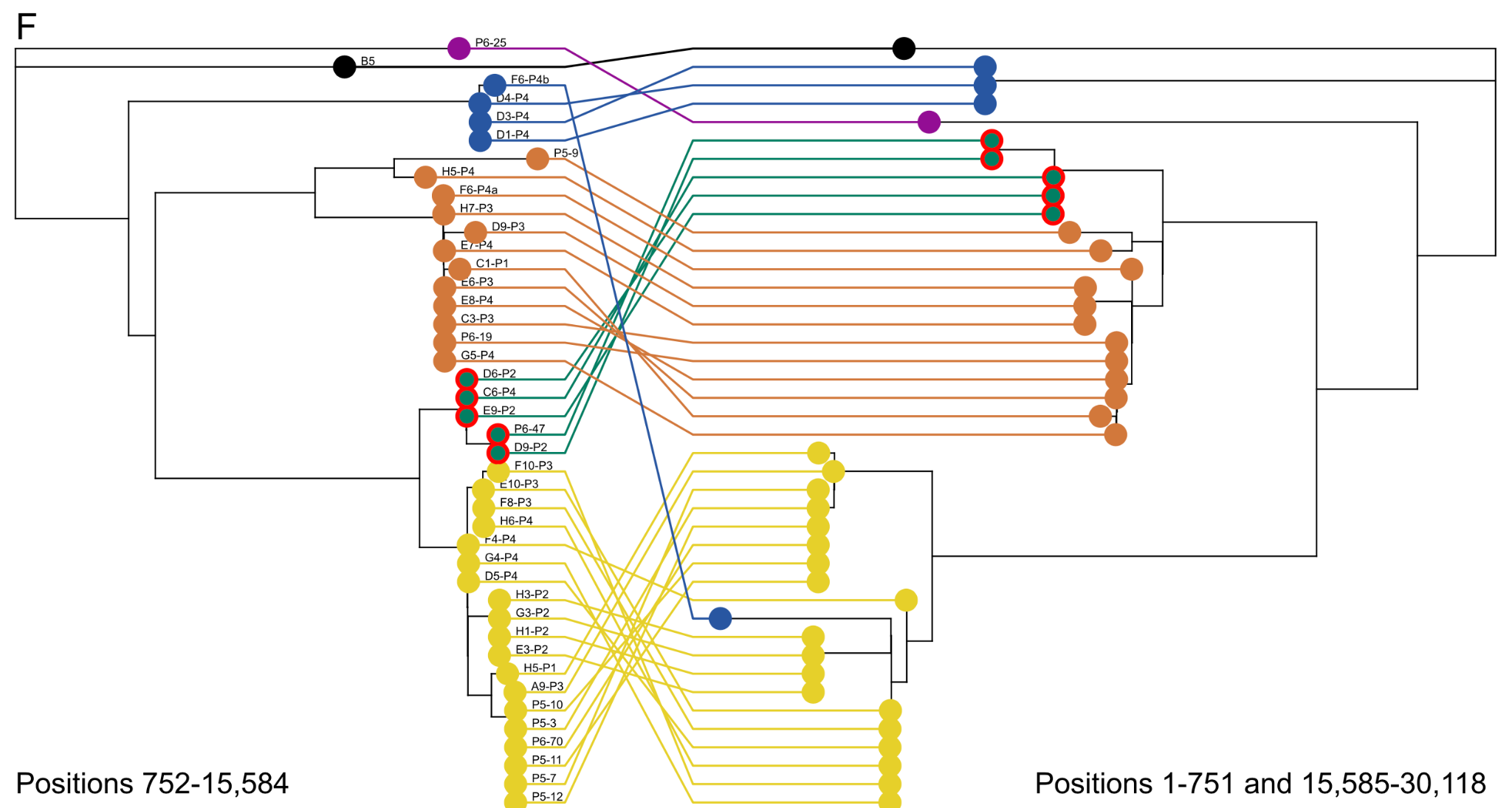

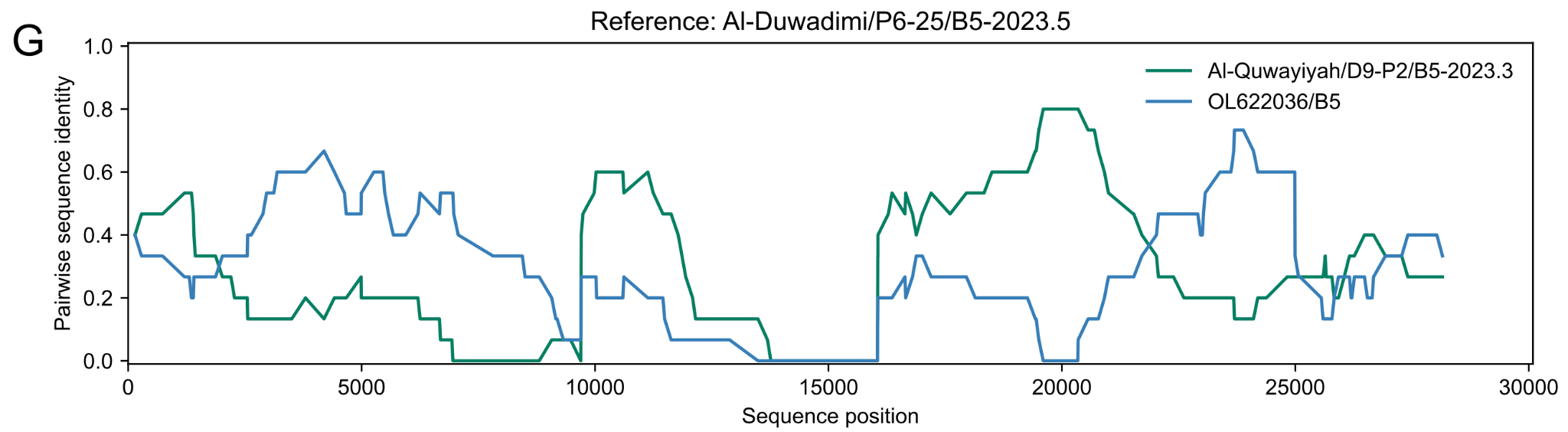

**Figure S2: Recombination analysis. A, B)** Plots showing a previously described recombination event between lineages B3, B4, and B5.[^26^](https://app.readcube.com/library/fefdf68c-7648-4f43-ae8d-20ef8d719102/all?uuid=7746086783897377&item_ids=fefdf68c-7648-4f43-ae8d-20ef8d719102:863ad90b-6246-4b3c-a27e-f5ec30cfbf3c) Pairwise sequence identity plots for lineages B3, B4, and B5 (A), and B3, B4, and a novel sequence (B). **C, D)** Sequence ‘Al Quwayiyah/F6-P4b/B5-2023.1’ results from a recombination event between sequences from the B5-2023.1 clade (blue) and the B5-2023.4 clade (yellow). C) Pairwise sequence identity plot between sequence ‘Al Quwayiyah/F6-P4b/B5-2023.1’ and a sequence from the B5-2023.1 clade (Sajir/D1-P4/B5-2023.1) and the B5-2023.4 clade (Al Quwayiyah/E3-P2/B5-2023.4). D) Trees made from the major (major: 1-17,816 and 29,588-30,118) and minor (offsets: 17817-29,587) parents. **E, F)** Sequences in the B5-2023.3 clade arose from a recombination event between sequences from the B5-2023.2 clade and sequences from the B5-2023.4 clade. E) Pairwise sequence identity plot between a sequence from the B5-2023.3 clade (Al Quwayiyah/D9-P2/B5-2023.3) and a sequence from the B5-2023.2 clade (Jeddah/P5-9/B5-2023.2) and the B5-2023.4 clade (Al Quwayiyah/E3-P2/B5-2023.4). F) Trees made from the major (offsets: 1-751 and 15,585-30,118) and minor (offsets: 752-15,584) parents. **G)** Pairwise sequence identity plot indicating a possible recombination event involving sequence Al Duwadimi/P6-25/B5-2023.5, between a sequence from clade B5-2023.3 (Al-Quwayiyah/D9-P2/B5-2023.3) and older sequences of lineage B5, represented here by OL622036/B5. Pairwise sequence identity shown in panels A, B and C and E was calculated excluding invariant sites and using a window size of 15 and a step size of 1. Panels D and F show trees inferred from the parents, with lines connecting the same sequence in the two trees. Recombinant sequences are indicated by a red outline.

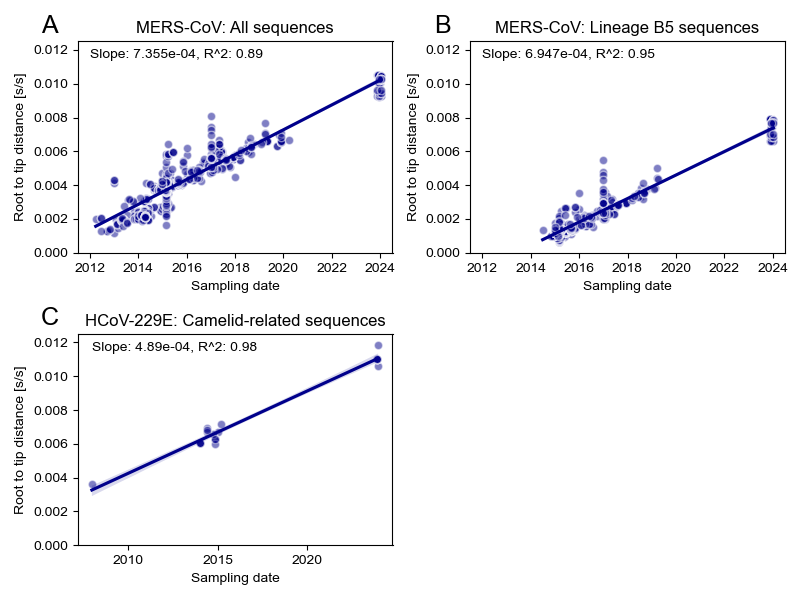

**Figure S3: Regression of root to top distances versus sampling date.** Root to tip distances are plotted against sampling dates for the full tree shown in figure 1A (A), for lineage B5 and the new clade (B), and for the camelid 229E-related sequences (C). Sampling dates where either the month and/or day were unknown, were set to the first of the month, or the first of January.

### **Tables**

**Table S1: Metadata for all tested samples.** Samples with identical IDs correspond to separate nasal swabs taken from the same camel.

Table will be provided as additional data.

**Table S2: Clade-defining substitutions for the subclades within the B5-2023 clade, based on the complete genome tree.**

|  | **B5-2023** | **B5-2023.1** | **B5-2023.2** | **B5-2023.3** | **B5-2023.4** | **B5-2023.5** |
| --- | --- | --- | --- | --- | --- | --- |
| **ORF1ab** | A1977T, D3889E, T642I, L759F | S611N, N682K, V698A, V1034I, N1147S, I2265T, T782I, T1740M | Y1888H, T1933I | N581S, T1740M | Q512H, D671N | S393N, S2138N, F2513L, Y2631F, R2961K, R1573G |
| **S** | R1179del, I1180del, L26V, A29S, S191P, A205S | K27E, S459N, H486Y, G1250V | None | R505L, A1206S, T1216I | P97L, G198D, L495P, V527L | V527L |
| **ORF3** | I44V, C38F, L86F | None | None | None | None | S101L |
| **ORF4b** | None | None | None | None | None | L41F, K151N |
| **ORF5** | V191A, I147T | None | None | None | None | None |
| **N** | N22H | None | None | G189C | None | A5S |
| **ORF8b** | None | None | None | None | None | Q68L, P4L |
| **Nucleotide** | G6207A, T11783G, C11864T, C11945A, T12092C, T13492C, C16363T, T16654C, C16879T, G18502T, C19237T, C20348A, G20695A, T20911C, C21531G, G21540T, C22391T, G24197A, A25661G, T26677C, T27411C, A28629C, T29165C, C2203T, C2553T, T11249C, C16801T, G19432A, C20272T, C20782T, T20998C, T22026C, G22068T, C24383T, G25644T, C25787T, T27279C, C28153T, C28257T, T11249C, T20911C, C21531G | C162T, G542T, T800C, T983C, G2110A, T2324G, T2371C, G3378A, A3718G, G6995A, T7072C, T7883C, C11666T, C13220T, T14903C, T16190G, T16225C, T16906C, C1346T, C2623T, C3419T, C4775T, C5497T, G7094A, C10301T, C11423T, G12761T, C16477T | T3078C, C5552T, T5940C, C6779T, C9656T, C12359T, T797C, C5162T, C6076T, C11423T, G17956T, A19449G, C21713T, C23012T, C21713T | G148A, G12884T, G22969T, G25071T, T27805C, G29130T, A2020G, C5497T, G17956T, A19449G, C21713T, C23012T, C25102T, C21713T | G1814T, G2289A, G6242A, G16915T, G17735A, C19264T, C19507T, C22913T, C23387T, C23447T, A17899T, C17959T, C18145T, T18502G, C18526T, C18949T, T19237C, T19313C, A20348C, G20522T, A20695G, T20782C, C20998T, T21055C, C21745T, G22048A, T22939C, G23034C, T24938C, C27073T, C18145T, T18502G, T19237C, C19264T, C19507T, A20348C, A20695G, T20782C, C20998T, C22913T, C23387T, C23447T | C287T, G1456A, T4673C, G6691A, G6806T, T7815C, A8170T, T8174C, G9160A, A10034G, C12155T, T13768C, C14144T, C15457T, G20341A, C20599A, T23072C, C23624T, T24920C, C26049T, T26197C, G26266A, C26932T, G28578T, G28718A, A28964T, T29441C, C1439T, C1778T, T2489C, T2639C, T2888A, C3506T, A4995G, C5948T, C9740T, A9983G, C10085T, G12518A, G12761T, G15535T, C16882T, C17606T, C18145T, T18334C, C21713T, G23034C, T25597C, C25833T, C25926T, C26213T, G26545T, C26647T, C28410T, C28772T, C9740T, C18145T, C21713T, T25597C, C28772T |

**Table S3: Clade defining substitutions for clades A and B, and different lineages within clade B.**

|  | **Clade A vs Clade B** | **Lineage B1-B4 vs lineage B5** | **Lineage B5 vs lineage B5-2023** |
| --- | --- | --- | --- |
| **ORF1ab** | K726N, P1055S, M1370I, L1717I, A588T, T1000I, R1573K, A2780V | None | A1977T, D3889E, T642I, L759F |
| **S** | Q1020H, Y194H, S1158A | None | S191P |
| **ORF3** | None | None | I44V, C38F, L86F |
| **ORF5** | None | None | V191A, I147T |
| **N** | None | None | N22H |
| **ORF8b** | L4P | None | None |
| **Nucleotide** | A2456T, C3134T, T3320C, C3441T, G4034A, G4388T, T5427A, A5516G, C6332T, C8258T, C10505T, T11984C, C12684T, G12707A, C13022T, C17794T, T26806C, G542A, C624T, C1514T, G2040A, C2318T, C3277T, G4996A, C8207T, C8333T, C8617T, G10835T, C11492T, A11534G, G16900T, G19075A, C20848A, T22035C, C22790T, T24299C, G24515C, G24704A, T24927G, C26716T, T28772C, T29811C | C23648T, C23756T, C23804T, G27067T, C27355T | G6207A, T11783G, C11864T, C11945A, T12092C, T13492C, C16363T, T16654C, C16879T, G18502T, C19237T, C20348A, G20695A, T20911C, C21531G, G21540T, C22391T, G24197A, A25661G, T26677C, T27411C, A28629C, T29165C, C2203T, C2553T, T11249C, C16801T, G19432A, C20272T, C20782T, T20998C, T22026C, G22068T, C24383T, G25644T, C25787T, T27279C, C28153T, C28257T |
